# Bird communities in a selectively logged tropical montane forest are dominated by small, low-elevation species

**DOI:** 10.1101/2023.04.15.537021

**Authors:** Ritobroto Chanda, Shambu Rai, Bharat Tamang, Binod Munda, D.K. Pradhan, Mangal Rai, Aman Biswakarma, Umesh Srinivasan

## Abstract

Tropical montane forests are critical centres of terrestrial biodiversity and endemism, and face a range of threats, including selective logging and climate change. However, few studies have explored the joint influence of these threats, particularly in tropical mountains, despite how crucial it is to understand the cumulative impact of forest loss and climate change on biodiversity. We used mist-netting and bird ringing data from a long-term (10-year) community and population monitoring program to examine how the composition of the mid-elevation Eastern Himalayan understorey bird community changed in the mid-elevation primary and logged forest. Because logged forest is warmer than primary forest, we hypothesised that the bird community would shift towards lower elevation species, with a faster transition in logged forest. Further, because logged forest has lower arthropod abundance, we hypothesised that the primary forest community would have larger species than the logged forest community. Our study shows that the bird community in the logged forest shifted towards lower-elevation species than in the primary forest (although this was not statistically significant). Moreover, we found that smaller species were better able to colonise warmer logged forest, whereas larger species reached higher densities in primary forest. These trends are likely to be driven by climate change causing upslope range shifts and logged forest selecting for species that are likely to tolerate the consequently altered abiotic (higher temperatures) and biotic (lower resources) environment. These findings have significant implications for understanding the impacts of forest loss and climate change on biodiversity, particularly in tropical montane forests.

## Introduction

Tropical montane forests are the richest centres of terrestrial biodiversity and endemism globally (Rahbek et al., 2019). These forests face multiple threats (Bradshaw et al. 2009), of which climate change and habitat degradation/conversion are major drivers of biodiversity declines (Asner et al., 2009; Edwards and Laurance, 2013; Constantini, Edwards, & Simons, 2016; Burivalova et al., 2014; Dawson et al., 2011; Corlett, 2012).

While the separate impacts of habitat loss and climate change on biodiversity have been studied extensively (Bellard et al., 2012; Matthews et al., 2014; Fahrig, 2003), few studies have examined the influence of both these threats together (Manytka-Pringle et al., 2012; Guo et al., 2018; Srinivasan and Wilcove, 2021), especially in the tropics, and particularly in tropical mountains. With climate change and forest loss accelerating dramatically (Mitchard, 2018; Arneth et al., 2020), understanding their cumulative impacts on biodiversity is essential. This is especially true because forest degradation (e.g., from selective logging) alters forest structure and creates novel habitats that tend to be hotter and drier than primary forest (Srinivasan and Wilcove, 2021), complicating climate impacts. Further, tropical species are thermal specialists and poor dispersers and tend to have narrow elevational ranges, and how climate- *and* land-use-driven changes in the abiotic environment might impact such species remains poorly known (Janzen, 1967; McCain, 2009; Sheard et al., 2020).

Tropical montane species are shifting their ranges to higher elevations rapidly, with strong evidence that such upslope shifts are a result of rising temperatures globally (Forero-Medina et al., 2011; Campos-Cerqueira et al., 2017; Freeman et al., 2018; Neate-Clegg et al., 2018, Freeman et al. 2021). As tropical species shift their ranges upslope in response to climate change, populations might either encounter primary forest or degraded forest (e.g., selectively logged forest) at higher elevations. With populations of the same species likely shifting their ranges into primary and logged forest at higher elevations, it is crucial to understand how such range shifts might lead to differences in community assembly in undisturbed versus disturbed forest. Although selectively logged forest can retain more biodiversity than other forms of forest loss/degradation such as outright conversion to agriculture (Putz et al., 2012; Gibson et al., 2011), logging does lead to the altered composition of ecological communities (Constantini, Edwards, & Simons, 2016; Burivalova et al., 2014).

Using mist-netting and bird ringing data from a long-term community and population monitoring program, we asked how the composition of the mid-elevation Eastern Himalayan understorey bird community changed over a ten-year period in primary and in logged forest. At any given elevation, the primary forest bird community should become progressively dominated by historically “low-elevation” species as they move upwards to replace “higher-elevation” species that are moving further upslope. Logged forest being warmer than primary forest (Srinivasan and Wilcove, 2021) should be thermally more similar to the lower-elevation primary forest than to the primary forest at the same elevation. We, therefore, expected that upslope range shifts should lead to faster species turnover from “higher-elevation” to “lower-elevation” in logged than in primary forest. Further, we expected that smaller species, being more thermally tolerant (Sheridan and Bickford, 2011; Speakman et al., 2003) would be better able to colonize warmer logged forest, whereas larger species would reach higher densities in primary forest.

We predicted that:

1. The bird community would gradually shift towards lower elevation species and the transition would occur faster in the logged forest.
2. The bird community in primary forest would have larger species as compared to the logged forest community.

## Methods

### Field Methods

We carried out this study in mid-elevation (2000m ASL) montane broadleaved forest (Champion and Seth, 1968) in Eaglenest Wildlife Sanctuary (27°06′0″N 92°24′0″E, EWLS henceforth), Arunachal Pradesh, Eastern Himalaya, India. In this habitat, we established three plots each in primary and logged forest (a total of 9ha each in primary and logged forest; Fig. 1). We selected the plots based on interviews with past logging managers. Intensive logging occurred until 2002 after which it was halted (see Srinivasan 2013). The logged patches have visible differences from the intact forest with a very open canopy and the understorey dominated by bamboo. We sampled each of these plots for three consecutive days from 0500 hrs to 1200 hrs in the early breeding season (April-May) from 2011 to 2021 using 24-28 mist nets (12m length, 2.4m height, 16mm mesh, 4-shelf). In 2020, a nationwide COVID-19-related lockdown precluded any bird sampling. We placed the mist nets in a grid-like pattern within the plot with ∼40m distance between neighbouring nets (Fig. 1). We then checked each net every 20-30 minutes and any birds captured were brought back to the ringing station at the centre of the sampling plot. We identified and weighed each bird caught with a jeweller’s scale, ringed it with a uniquely numbered aluminium ring and then released the bird immediately.

**Figure 1:**
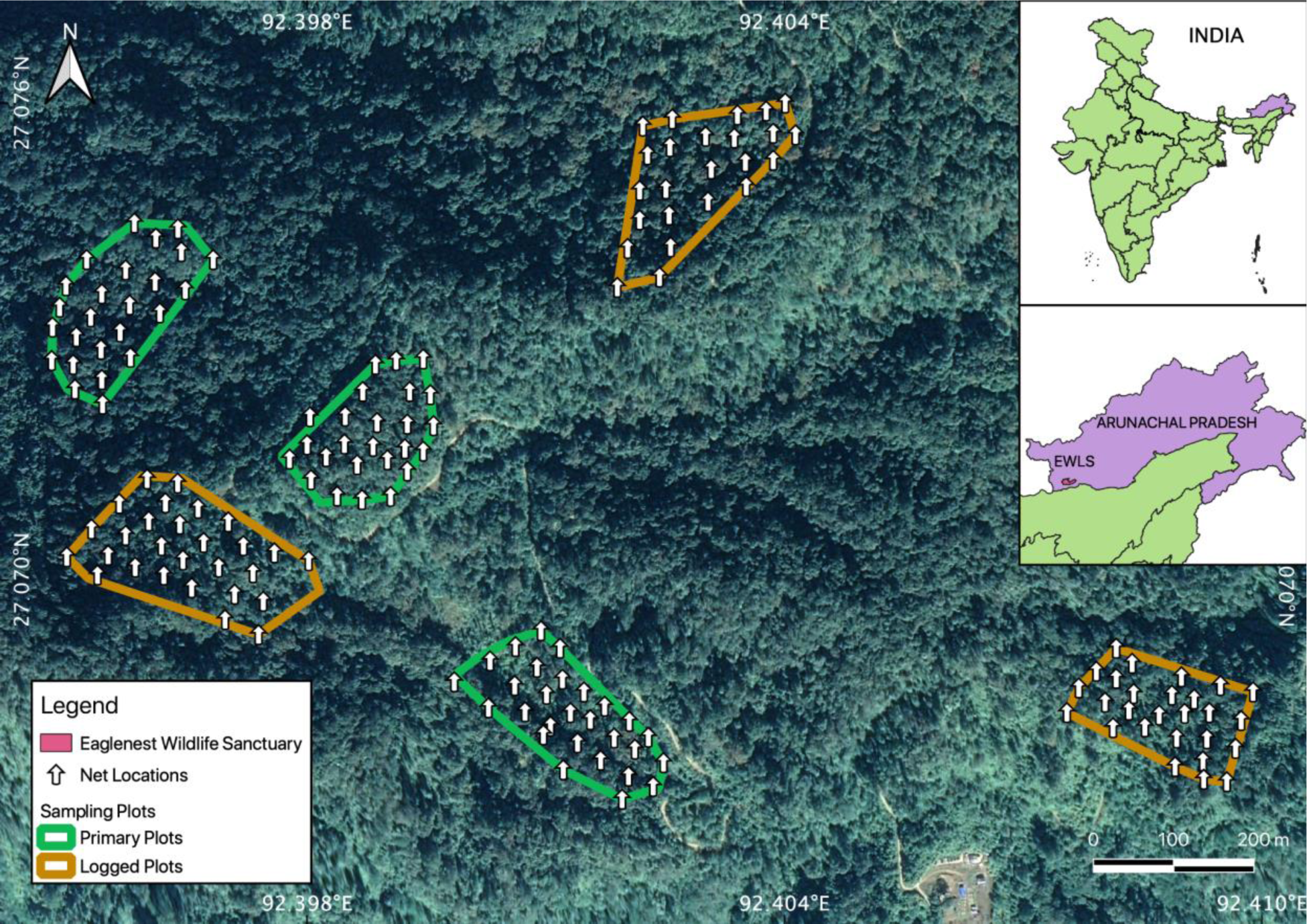
Map of the study area in Eaglenest Wildlife Sanctuary, Arunachal Pradesh, India. Primary forest and logged forest plots are outlined in green and brown, respectively. Arrows show the locations of mist nets.

### Data Curation

We excluded high-elevation breeders which are passage migrants to higher-elevation breeding sites and do not breed in our study area (Table S1), aquatic species, carnivores, canopy dwellers, and non-invertivores (which included frugivores, granivores and nectarivores). We finally analysed the understorey insectivorous bird community of 61 species (Table S1). We pooled the raw counts of each species in each habitat in each year pooled from for the first two days of sampling effort in each site (i.e., six days each in logged and primary forest in each year). We did this because in some years in some plots, rain prevented the full three-day sampling protocol and sampling was limited to two consecutive days.

### Analytical Methods

#### Differences in community composition

We first created a year-by-species matrix in primary forest and logged forest separately with the raw count data (Table S2) transformed using the Hellinger transformation (Bocard et al., 2011) which reduces the weightage of rare species in contributing to compositional differences between communities (Legendre and Gallagher, 2001). We then performed a non-metric multidimensional scaling (NMDS) ordination analysis to visualise the bird community in logged and primary forests each year using the Bray-Curtis dissimilarity index on the transformed data in the R package *vegan* (Okansen et al., 2022; R Core Team 2023). To test whether the bird community composition differed significantly between primary and logged forest, we used permutational multivariate analysis of variance (PERMANOVA), again using the Bray-Curtis dissimilarity index.

To understand the relative contributions of different species to community-level differences between primary and logged forest, we used similarity percentage routines (SIMPER, Clarke and Warwick 2001). From the SIMPER routine, we then selected the top species that cumulatively contributed to 50% of dissimilarity in bird community composition between primary and logged forest (Table S3). To visualise the importance of these species to the difference in community composition across habitats, we plotted the location of these species in NMDS space. We used SIMPER since it gives us a much clearer picture of the possible patterns than using all of the 61 species, especially rarer species that might have had only a few individuals captured over the study period and might skew the results disproportionately, appearing to contribute much to dissimilarity between communities, but not indeed doing so.

To understand potential differences between the morphology (i.e., body mass) and distribution (i.e., elevational ranges) of species that contributed in large part to community-level differences between primary and logged forest, we used:

a. A generalized linear model (GLMs) with habitat type (logged/primary) as the categorical predictor variable and body mass as the response variable.
b. An ANOVA to compare the elevation range midpoints of species in primary and logged forest.

The body mass of the species was averaged from the birds caught in logged and primary forests over the ten years. We extracted the breeding elevation range data of each from Spierenburg (2005) and it was supplemented by data from Srinivasan et al. (2018). We used this data to calculate the elevation range midpoint.

#### Trends over time

To understand the trends of body mass and elevation range midpoints over time, we came up with two metrics, community mass index (CMI) and community elevation index (CEI). These were developed using a metric called the community temperature index (Devictor et al., 2008). We took the community weighted mean of species mass and the breeding elevation range midpoint to understand the temporal shift. We carried this out using the top species that cumulatively contributed to 50% of dissimilarity in bird community composition between primary and logged forest.

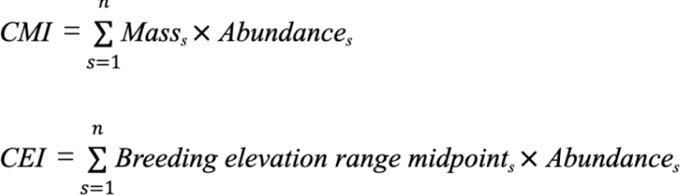

## Results

We recorded 9,036 captures of 6,189 individuals from 130 species in our mist nets across all years and across all sampling plots. Upon curating the data, we finally analysed 6,926 captures of 4,801 individuals of 61 understorey insectivorous species. The raw counts of the 61 species are in Table S2.

### Community composition in primary and logged forest

NMDS axis 1 separated bird communities in primary and logged habitats (stress = 0.20, Fig. 2). Primary and logged forest communities were compositionally significantly different from each other (PERMANOVA (R^2^ = 0.14, F_1,18_ = 2.98, *p* < 0.01). The first 20 species (Table S3) explained 50% of the variation in community composition between primary and logged forest and were used in subsequent analyses. We did not use *Schoeniparus cinereus* and *Tesia castaneocoronata* out of these since they did not fall in either primary or logged forest clusters.

**Figure 2:**
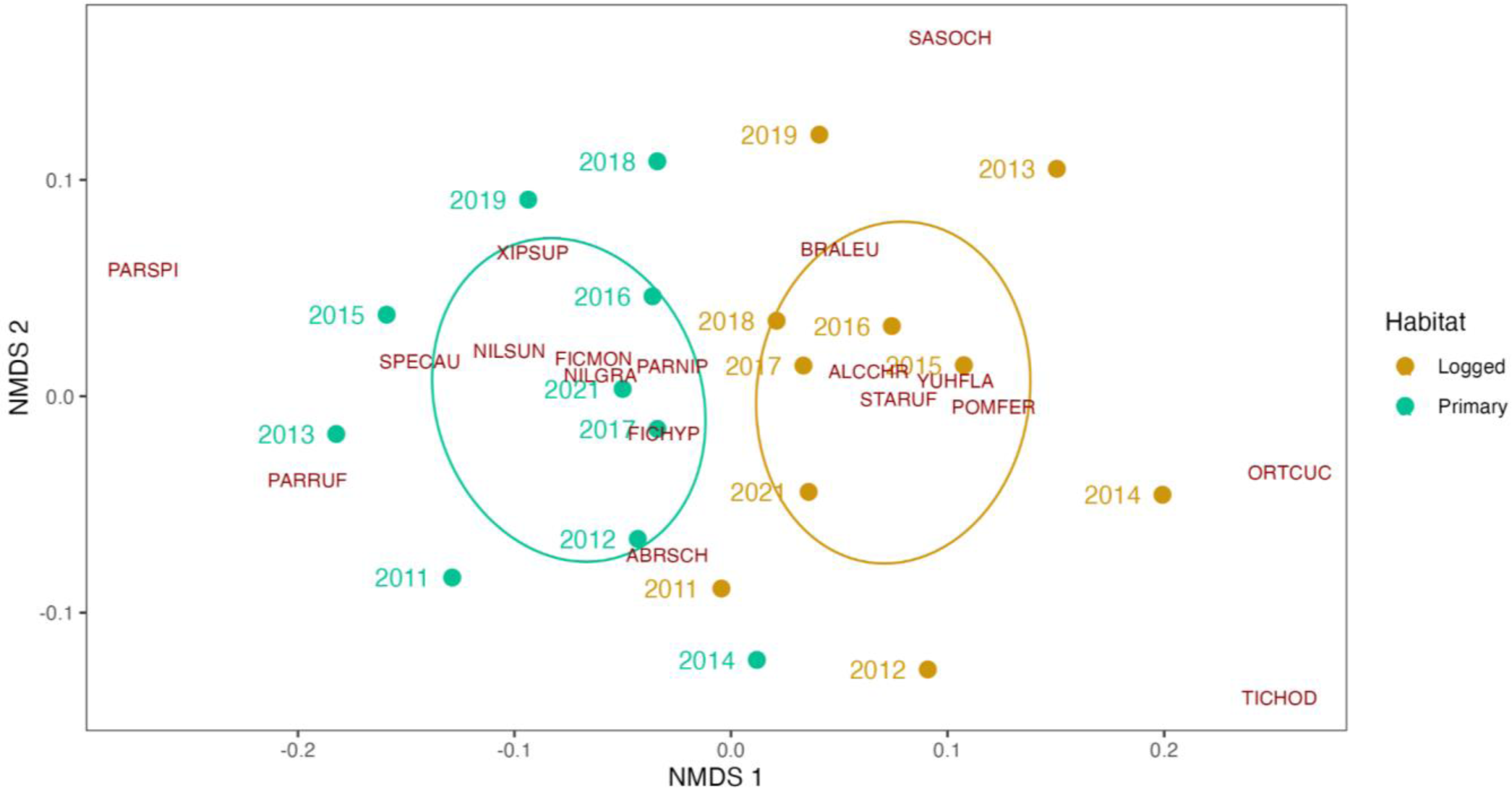
Non-metric multidimensional scaling (NMDS) showing dissimilarity between primary and logged forest communities over a ten-year period (see Table S3 for details of species codes and contribution to the SIMPER routine).

### Body mass trends in primary and logged forest

Habitats were significantly related to body mass (Poisson ANOVA, ***β***_primary_ = 0.26 [0.02, 0.50], McFadden’s Pseudo *R^2^* = 0.04. On average, species typical to primary forest were 1.3g heavier than species typical to logged forest. Median body mass across species was 5g higher in primary forest (Fig. 3).

**Figure 3:**
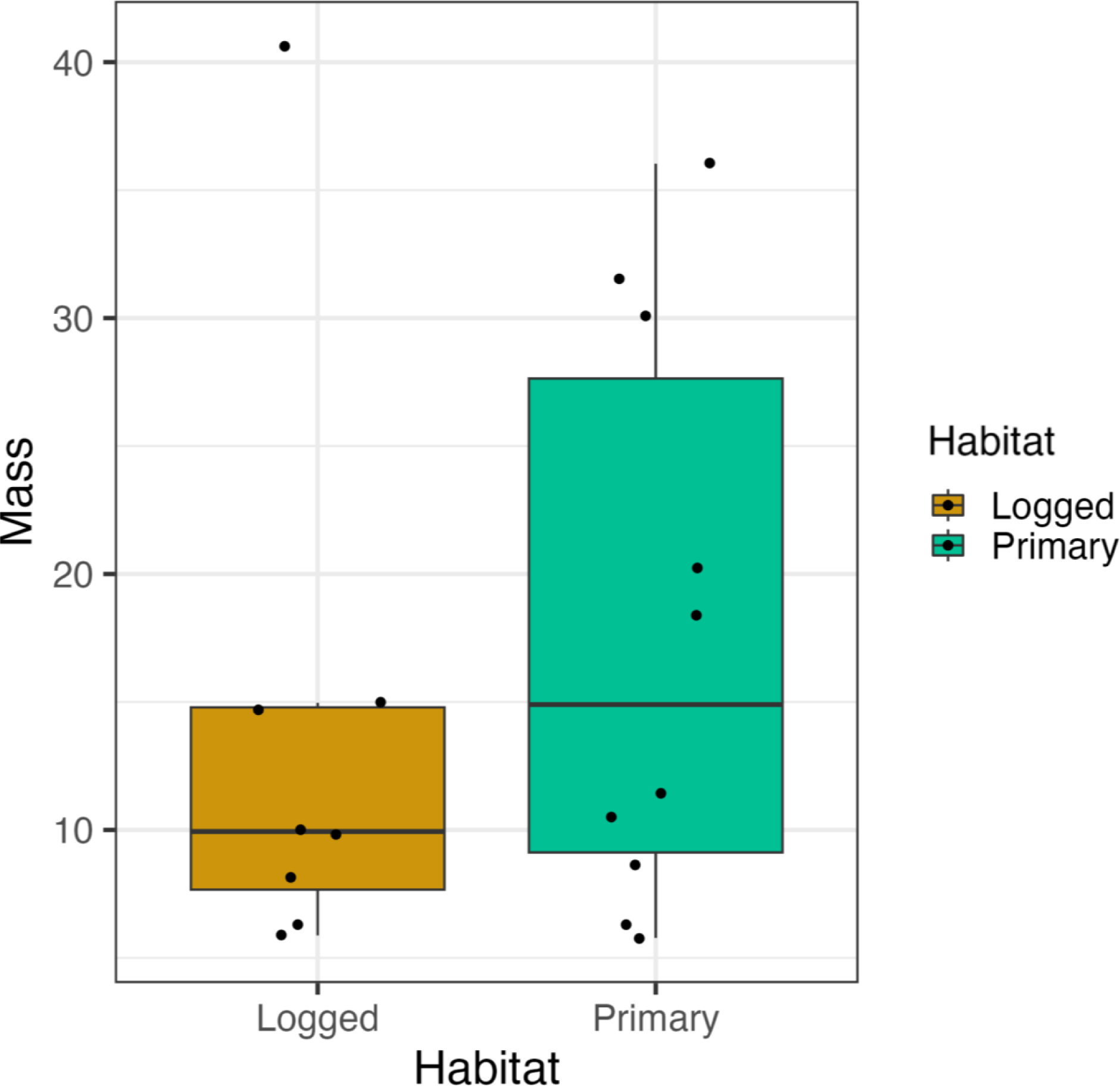
Mass of species that contributed to 50% of dissimilarity in community composition between primary and logged forests. Species typical to primary forest are, on average, larger than those typical to logged forest. The box shows the middle 50% of the data, with the horizontal line inside the box indicating the median value. The whiskers of the boxplot extend from the box to show the minimum and maximum values of the data. Each dot represents individual data points and is plotted with a jitter to avoid overlap.

### Trends of Breeding elevational range midpoints in primary and logged forest

Elevation range midpoints of bird species typical to primary forest was, on average 172.5m [-185.3, 530.3; *R^2^* = 0.06] higher than species representative of logged forest. While the mean difference in the mid-points of species’ elevational ranges was as expected, the effect of habitat was weak (i.e., 95% CIs overlapping zero).

The mean low elevation range limits of species in the primary forest were 175m [-188.9, 538.9; *R^2^* = 0.06] higher than in logged forest. Similarly, the mean high elevation range limit of the species was higher in primary forest as well by 170 m [-270.3, 610.3; *R^2^* = 0.04], supporting the overall trend (although weak) of species representative of primary forest being “higher elevation” species than those representative of logged forest in the mid-elevation montane temperate forest (See Supp. Figs. S1 and S2).

### Trends over time

We tried to look at the trends of mass and mid-point of breeding elevation range over time. We found major overlaps in the confidence intervals of CEI and CMI over time (see Supp. Figs. S3 and S4).

## Discussion

We investigated how climate change and selective logging might combine to impact tropical understorey bird communities in primary and selectively logged forest using a 10-year dataset from the Eastern Himalayas. We find that the bird community in primary forest was, on average, composed of larger species than in logged forest (Fig. 3). We also find weak support that the community in logged forest has a lower mean, minimum and maximum summer elevation range than the primary forest community, which indicates that the logged forest community might resemble lower elevation communities (Fig. 4). However, this effect was not as strong as the effect for body mass. We tried to understand trends of mass and mid-point of breeding elevation range over time but this seemed to be complicated by natural trends in abundance fluctuations in each year (Supp. Fig S3 and S4). We can likely see stronger trends with more long-term data.

**Figure 4:**
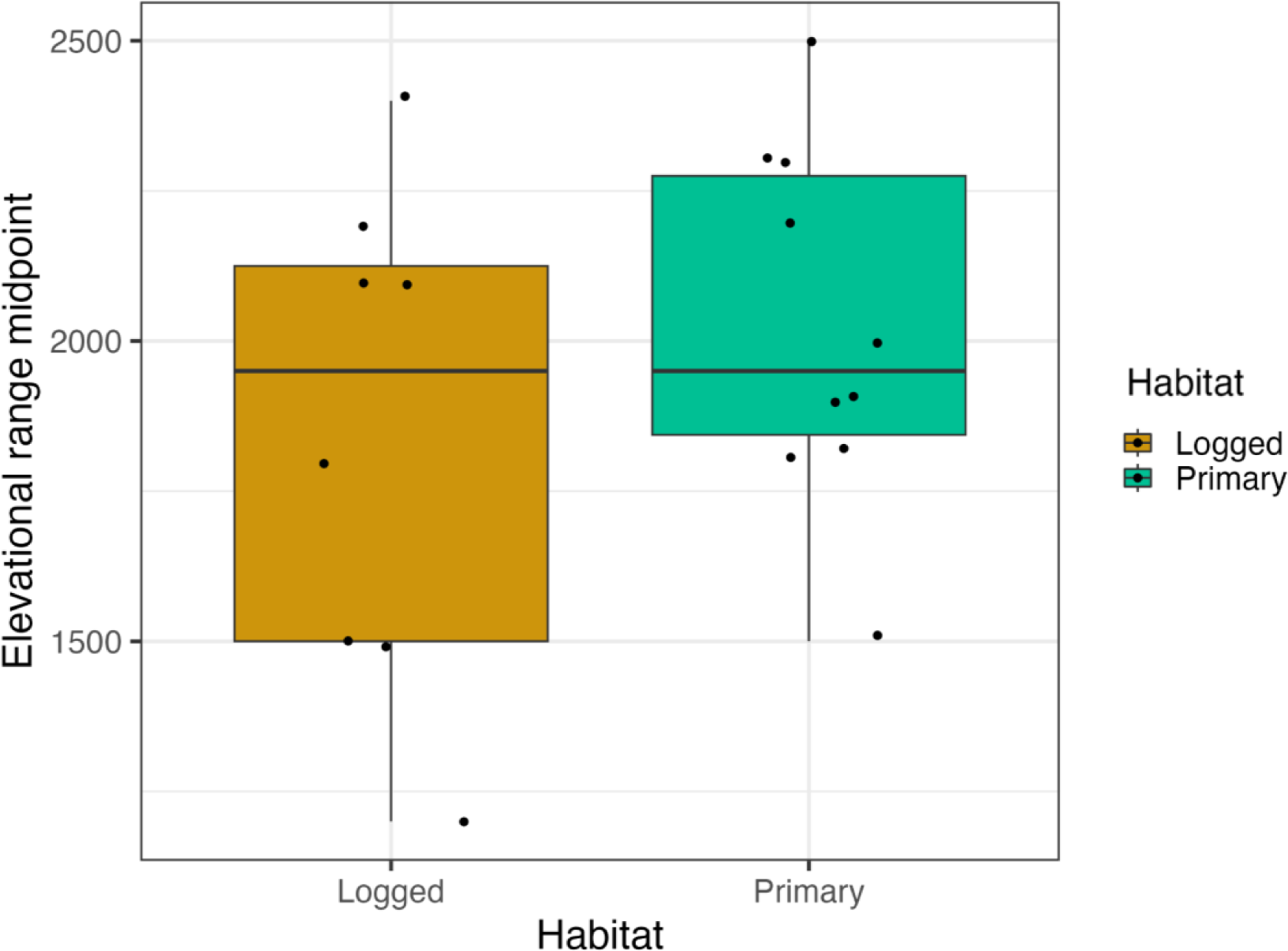
Mean breeding elevation range of the species that contributed to 50% dissimilarity in each habitat. The box shows the middle 50% of the data, with the horizontal line inside the box indicating the median value. The whiskers of the boxplot extend from the box to show the minimum and maximum values of the data. Each dot represents individual data points and is plotted with a jitter to avoid overlap.

Logging can lead to the loss of large-bodied, old-growth-dependent species and decrease overall biodiversity (Cleary et al., 2007). Understorey insectivores, which are often highly specialized, are negatively influenced by logging and show steep abundance declines (Chaves et al. 2017). Further, the body size of birds is an important factor affecting their response to logging. Birds with larger body sizes were found to be less common in logged forests, whereas smaller-bodied species were found to be more resilient to logging (Burivalova et al., 2015). We find a similar pattern, with species typical of primary forest being significantly larger than species typical of logged forest (Fig. 2). Smaller-bodied bird species were more abundant in logged forest, while larger-bodied species declined (Velho et al., 2012, Srinivasan 2013). Such patterns might arise from changes in the abiotic environment with logging leading to changes in arthropod prey composition and abundance, in turn resulting in reduced resource availability for birds (Aggarwal et al., 2023). Earlier studies from our field site have shown that logged forest is warmer than old-growth forest because of changes in forest structure and microclimates (Bharadwaj et al., 2022). On average, primary forest was found to be 2.3°C cooler and 14.6% more humid than logged forest (Aggarwal et al., 2023), and have lower densities of foliage-dwelling arthropods (Aggarwal et al., 2023). Because large species have higher absolute energy requirements, a reduction in resource availability is likely to disproportionately reduce the abundance of large species (Srinivasan 2013).

The altered abiotic environment in logged forest can not only reduce the abundance of large species through indirect impacts on prey availability but can also directly influence the composition of bird communities in terms of body mass and elevational range. Climate warming is believed to have driven a decrease in the body size of warm-blooded animals and recent work in the Neotropics has found mean body mass and the mass: wing length ratio of birds decreasing over the past four decades (Jirinec et al., 2021). Smaller individuals and species might do better in warmer habitats for several reasons. Smaller individuals tend to have a higher surface area to volume ratio, which means they lose heat more quickly and can adapt to warmer temperatures more easily (Sheridan and Bickford, 2011; Speakman et al., 2003). Further, smaller organisms tend to have higher metabolic rates, which enables them to grow and reproduce faster in warmer environments (Angilletta, 2009).

Across the world’s tropical mountains, bird populations are shifting to higher elevations in response to rising temperatures (Forero-Medina et al., 2011; Freeman et al., 2018; Neate-Clegg et al., 2018; Freeman and Class Freeman, 2014), including in the Eastern Himalaya (Srinivasan & Wilcove, 2021; Girish and Srinivasan, 2022). At our study site, as populations shift their elevational ranges upslope, they might encounter primary or logged forest. Our results show that logged forest bird communities are dominated by species that have a mean elevation range midpoint that is lower than that of primary forest species, although this effect was weak (Fig. 4). Logged forest, which is consistently warmer than primary forest at our site (with a mean midday difference of 2.3°C, Aggarwal et al., 2023) might be more suitable abiotically for lower-elevation, comparatively warm-adapted, species compared with primary forest.

These indirect impacts of climate change on bird species highlight the complex interplay between biotic and abiotic factors in shaping ecological communities. The weak shift in species composition of understory birds in logged forest towards communities resembling those at lower elevations suggests that while climate change will cause upslope range shifts, habitat degradation might then select for species that are adapted to the altered abiotic conditions typical of such habitats. At our site, bird populations at the cold-edge elevational range limit have increased survival rates over time, while those at the warm-edge elevational range limit suffer declines (Srinivasan and Wilcove, 2021). This suggests that species are adapting to climate change by tracking favourable conditions and shifting their ranges upslope. However, individuals of the same species are unable to maintain demographic vital rates in higher-elevation logged forests in response to climate change in the long term (Srinivasan and Wilcove, 2021). This could be due to a decrease in resource availability in logged forest which can severely impact bird demographics (Grames et al., 2023). Therefore, protecting large tracts of well-preserved primary forests across elevational gradients is essential for enabling tropical montane species to survive the dual effects of climate change and land-use change.

This study aimed to investigate the effects of climate change and selective logging on the insectivorous bird communities in tropical understorey in primary and selectively logged forests in the Eastern Himalayas of India. The results indicate that smaller species, more typical of low elevations, tended to dominate the bird communities in logged forests. This shift in body size and range may be due to various factors, such as changes in food resources, habitat, and microclimate resulting from logging and warming temperatures. This emphasises the importance of considering long-term changes in bird communities when developing conservation strategies for bird species and their habitats. In light of our findings, continuous monitoring and management of bird habitats are crucial to prevent declines in bird diversity and abundance.

## Acknowledgements

We thank the Arunachal Pradesh Forest Department for their continuous support and for providing us with the permits to carry out this work. We are thankful to the Indian Institute of Science, Science and Engineering Research Board, GoI, and the Department of Biotechnology, GoI for funding our study. We thank Indi Glow, Nima Tsering, Joli Borah, Ramesh Kumar Pradhan, Kushal Hazarika, LN Thongchi, Millo Tasser, Nana Khrimey, Pema Mosobi, Ramana Athreya, Puspalal Sharma, Dorjee Pema, Satu Narah, Shreesh Kaulgud and Tarun Menon for all their help during fieldwork. We are grateful towards the people of Ramalingam, West Kameng for their constant support.

## Ethics statement

We thank the Arunachal Pradesh Forest Department for providing us with permission for this study (No: CWL/GEN/2018-19/Pt.IX/NG/5210-13). This study has been approved by the Institutional Animal Ethics Committee of the Indian Institute of Science (No: CAF/Ethics/883/2022).

## Supplementary materials

**Supp. Table S1:** Final list of bird species used for analyses after excluding high-elevation breeders that do not breed in our study area, aquatic species, carnivores, canopy dwellers, and non-invertivores (which included frugivores, granivores and nectarivores).

|  | Scientific Name | Species Code | English Name | Mass (g) | Forest Layer | Diet | Migrant | Flocking | Elevation - Summer Low | Elevation - Summer High | Elevation - Summer Mean |
| --- | --- | --- | --- | --- | --- | --- | --- | --- | --- | --- | --- |
| 1 | *Abroscopus schisticeps* | ABRSCH | Black-faced Warbler | 5.78 | Midstorey | Insectivore | No | Yes | 1400 | 2400 | 1900 |
| 2 | *Actinodura egertoni* | ACTEGE | Rusty-fronted Barwing | 32.77 | Midstorey | Insectivore | No | Yes | 1000 | 2000 | 1500 |
| 3 | *Actinodura nipalensis* | ACTNIP | Hoary-throated Barwing | 46.46 | Midstorey | Insectivore | No | Yes | 1800 | 3200 | 2500 |
| 4 | *Actinodura waldeni* | ACTWAL | Streak-throated Barwing | 47.5 | Midstorey | Insectivore | No | Yes | 2400 | 3300 | 2850 |
| 5 | *Aegithalos concinnus* | AEGCON | Black-throated Bushtit | 6.14 | Midstorey | Insectivore | No | Yes | 1200 | 2600 | 1900 |
| 6 | *Alcippe castaneiceps* | ALCCAS | Rufous-winged Fulvetta | 9.7 | Midstorey | Insectivore | No | Yes | 1800 | 3200 | 2500 |
| 7 | *Alcippe cinerea* | ALCCIN | Yellow-throated Fulvetta | 9.34 | Undertorey | Insectivore | No | Yes | 1400 | 2400 | 1900 |
| 8 | *Anthipes monileger* | FICMON | White-gorgeted Flycatcher | 11.41 | Midstorey | Insectivore | Yes | No | 1600 | 2000 | 1800 |
| 9 | *Blythipicus pyrrhotis* | BLYPYR | Bay Woodpecker | 164.49 | Midstorey | Insectivore | Yes | Yes | 800 | 2200 | 1500 |
| 10 | *Brachypteryx leucophris* | BRALEU | Lesser Shortwing | 14.73 | Terrestrial | Insectivore | Yes | No | 1400 | 2800 | 2100 |
| 11 | *Brachypteryx montana* | BRAMON | White-browed Shortwing | 18.57 | Terrestrial | Insectivore | Yes | No | 2200 | 3400 | 2800 |
| 12 | *Certhia discolor* | CERDIS | Brown-throated Treecreeper | 10.86 | Midstorey | Insectivore | Yes | Yes | 160 | 2600 | 1380 |
| 13 | *Cettia fortipes* | CETFOR | Brown-flanked Bush Warbler | 10.23 | Undersorey | Insectivore | Yes | No | 1400 | 2800 | 2100 |
| 14 | *Cissa chinensis* | CISCHI | Common Green Magpie | 127 | Midstorey | Omnivore | No | No | 400 | 2000 | 1200 |
| 15 | *Dendrocitta frontalis* | DENFRO | Collared Treepie | 80.99 | Midstorey | Omnivore | No | No | 600 | 2000 | 1300 |
| 16 | *Ficedula hyperythra* | FICHYP | Snowy-browed Flycatcher | 8.65 | Undersorey | Insectivore | Yes | No | 1800 | 2800 | 2300 |
| 17 | *Ficedula westermanni* | FICWES | Little Pied Flycatcher | 7.8 | Midstorey | Insectivore | Yes | No | 1000 | 2400 | 1700 |
| 18 | *Garrulax caerulatus* | GARCAE | Grey-sided Laughingthrush | 80.61 | Undersorey | Omnivore | No | Yes | 1600 | 2400 | 2000 |
| 19 | *Harpactes erythrocephalus* | HARERY | Red-headed Trogon | 100.81 | Midstorey | Omnivore | Yes | Yes | 400 | 2000 | 1200 |
| 20 | *Hydrornis nipalensis* | PITNIP | Blue-naped Pitta | 124 | Terrestrial | Insectivore | Yes | No | 1400 | 1800 | 1600 |
| 21 | *Leiothrix argentea* | LEIARG | Silver-eared Leiothrix | 21.89 | Undersorey | Omnivore | No | No | 600 | 1800 | 1200 |
| 22 | *Leiothrix lutea* | LEILUT | Red-billed Leiothrix | 22.36 | Undersorey | Omnivore | No | No | 1800 | 3200 | 2500 |
| 23 | *Liocichla bugunorum* | LIOBUG | Bugun Liocichla | 39 | Midstorey | Omnivore | No | Yes | 2000 | 2600 | 2300 |
| 24 | *Lioparus chrysotis* | ALCCHR | Golden-breasted Fulvetta | 8.12 | Undersorey | Insectivore | No | Yes | 2000 | 2800 | 2400 |
| 25 | *Machlolophus spilonotus* | PARSPI | Yellow-cheeked Tit | 18.39 | Midstorey | Insectivore | Yes | Yes | 1200 | 2450 | 1825 |
| 26 | *Muscicapa ferruginea* | MUSFER | Ferruginous Flycatcher | 10.21 | Midstorey | Insectivore | Yes | No | 1400 | 2800 | 2100 |
| 27 | *Muscicapella hodgsoni* | MUSHOD | Pygmy Flycatcher | 5.76 | Midstorey | Insectivore | Yes | No | 1400 | 2600 | 2000 |
| 28 | *Myiomela leucura* | MYILEU | White-tailed Robin | 25.72 | Terrestrial | Insectivore | Yes | No | 1200 | 2200 | 1700 |
| 29 | *Niltava grandis* | NILGRA | Large Niltava | 36.03 | Midstorey | Insectivore | Yes | No | 1600 | 2200 | 1900 |
| 30 | *Niltava macgrigoriae* | NILMAC | Small Niltava | 11.31 | Understorey | Insectivore | Yes | No | 800 | 2000 | 1400 |
| 31 | *Niltava sundara* | NILSUN | Rufous-bellied Niltava | 20.2 | Understorey | Insectivore | Yes | No | 1600 | 3000 | 2300 |
| 32 | *Parus monticolus* | PARMON | Green-backed Tit | 18.65 | Midstorey | Insectivore | Yes | Yes | 1200 | 3000 | 2100 |
| 33 | *Phyllergates cuculatus* | ORTCUC | Mountain Tailorbird | 6.31 | Understorey | Insectivore | Yes | Yes | 1400 | 2200 | 1800 |
| 34 | *Phylloscopus chloronotus* | PHYCHL | Lemon-rumped Warbler | 5.01 | Midstorey | Insectivore | Yes | Yes | 2400 | 3600 | 3000 |
| 35 | *Phylloscopus reguloides* | PHYREG | Blyth's Leaf Warbler | 7.05 | Midstorey | Insectivore | Yes | Yes | 1600 | 3200 | 2400 |
| 36 | *Pnoepyga pusilla* | PNOPUS | Pygmy Wren-babbler | 12.25 | Understorey | Insectivore | No | Yes | 1000 | 2600 | 1800 |
| 37 | *Pomatorhinus ferruginosus* | POMFER | Coral-billed Scimitar Babbler | 40.65 | Understorey | Insectivore | No | Yes | 1000 | 2000 | 1500 |
| 38 | *Pomatorhinus ruficollis* | POMRUF | Streak-breasted Scimitar Babbler | 29.76 | Midstorey | Insectivore | Yes | Yes | 1600 | 2600 | 2100 |
| 39 | *Pomatorhinus superciliosus* | XIPSUP | Slender-billed Scimitar Babbler | 30.12 | Understorey | Omnivore | Yes | No | 1600 | 3400 | 2500 |
| 40 | *Psarisomus dalhousiae* | PSADAL | Long-tailed Broadbill | 55.06 | Midstorey | Insectivore | Yes | Yes | 400 | 2000 | 1200 |
| 41 | *Psittiparus ruficeps* | PARRUF | White-breasted Parrotbill | 31.54 | Midstorey | Insectivore | No | Yes | 1000 | 2000 | 1500 |
| 42 | *Rhipidura albicollis* | RHIALB | White-throated Fantail | 12.02 | Midstorey | Insectivore | Yes | Yes | 1200 | 2400 | 1800 |
| 43 | *Rimator malacoptilus* | RIMMAL | Long-billed Wren-Babbler | 19.69 | Undersorey | Insectivore | Yes | Yes | 900 | 2700 | 1800 |
| 44 | *Sasia ochracea* | SASOCH | White-browed Piculet | 10.03 | Midstorey | Insectivore | Yes | Yes | 400 | 2000 | 1200 |
| 45 | *Seicercus affinis* | SEIAFF | White-spectacled Warbler | 7.58 | Midstorey | Insectivore | Yes | Yes | 1000 | 2800 | 1900 |
| 46 | *Seicercus poliogenys* | SEIPOL | Grey-cheeked Warbler | 6.57 | Midstorey | Insectivore | Yes | Yes | 1400 | 2400 | 1900 |
| 47 | *Spelaornis caudatus* | SPECAU | Rufous-throated Wren-Babbler | 10.53 | Terrestrial | Insectivore | No | Yes | 1600 | 2400 | 2000 |
| 48 | *Spelaornis formosus* | ELAFOR | Spotted Wren-Babbler | 10.79 | Terrestrial | Insectivore | No | No | 800 | 2200 | 1500 |
| 49 | *Sphenocichla humei* | SPHHUM | Sikkim Wedge-billed Babbler | 34.1 | Undersorey | Insectivore | No | Yes | 1400 | 2000 | 1700 |
| 50 | *Stachyridopsis chrysaea* | STACHR | Golden Babbler | 7.85 | Undersorey | Insectivore | No | Yes | 800 | 2000 | 1400 |
| 51 | *Stachyridopsis ruficeps* | STARUF | Rufous-capped Babbler | 9.85 | Undersorey | Insectivore | Yes | Yes | 1200 | 1800 | 1500 |
| 52 | *Stachyris nigriceps* | STANIG | Grey-throated Babbler | 14.17 | Undersorey | Insectivore | No | Yes | 400 | 1800 | 1100 |
| 53 | *Suthora nipalensis* | PARNIP | Black-throated Parrotbill | 6.31 | Undersorey | Insectivore | No | Yes | 1600 | 2800 | 2200 |
| 54 | *Tesia castaneocoronata* | TESCAS | Chestnut-headed Tesia | 8.31 | Terrestrial | Insectivore | Yes | No | 2200 | 3400 | 2800 |
| 55 | *Tesia cyaniventer* | TESCYA | Grey-bellied Tesia | 9.89 | Terrestrial | Insectivore | Yes | No | 1400 | 2600 | 2000 |
| 56 | *Tesia olivea* | TESOLI | Slaty-bellied Tesia | 8.79 | Terrestrial | Insectivore | Yes | Yes | 1400 | 2000 | 1700 |
| 57 | *Tickellia hodgsoni* | TICHOD | Broad-billed Warbler | 5.88 | Understorey | Insectivore | Yes | Yes | 1600 | 2600 | 2100 |
| 58 | *Trochalopteron erythrocephalum* | GARERY | Chestnut-crowned Laughingthrush | 71.63 | Understorey | Omnivore | No | Yes | 1600 | 3400 | 2500 |
| 59 | *Trochalopteron squamatum* | GARSQU | Blue-winged Laughingthrush | 81.48 | Understorey | Omnivore | No | Yes | 1000 | 2000 | 1500 |
| 60 | *Trochalopteron subunicolor* | GARSUB | Scaly Laughingthrush | 66 | Understorey | Omnivore | No | Yes | 2000 | 3400 | 2700 |
| 61 | *Yuhina flavicollis* | YUHFLA | Whiskered Yuhina | 14.97 | Midstorey | Omnivore | Yes | No | 1600 | 2800 | 2200 |

**Supp. Table S2:** Raw counts of species in primary and logged forest. The first ten columns are from primary, followed by logged.

|  | Species Code | 2011.P | 2012.P | 2013.P | 2014.P | 2015.P | 2016.P | 2017.P | 2018.P | 2019.P | 2020.P | 2011.L | 2012.L | 2013.L | 2014.L | 2015.L | 2016.L | 2017.L | 2018.L | 2019.L | 2021.L |
| --- | --- | --- | --- | --- | --- | --- | --- | --- | --- | --- | --- | --- | --- | --- | --- | --- | --- | --- | --- | --- | --- |
| 1 | ABRSC H | 3 | 5 | 4 | 10 | 3 | 2 | 5 | 0 | 1 | 5 | 3 | 2 | 0 | 2 | 1 | 6 | 5 | 3 | 3 | 3 |
| 2 | ACTEG E | 4 | 9 | 13 | 4 | 8 | 4 | 6 | 2 | 4 | 6 | 1 | 7 | 5 | 6 | 6 | 13 | 6 | 14 | 5 | 10 |
| 3 | ACTNI P | 0 | 0 | 0 | 0 | 0 | 0 | 0 | 0 | 0 | 1 | 0 | 0 | 0 | 0 | 0 | 0 | 0 | 1 | 0 | 0 |
| 4 | ACTW AL | 1 | 0 | 2 | 0 | 0 | 1 | 0 | 0 | 0 | 0 | 0 | 2 | 0 | 0 | 1 | 1 | 0 | 0 | 0 | 0 |
| 5 | AEGCO N | 0 | 1 | 0 | 0 | 1 | 0 | 3 | 0 | 0 | 0 | 0 | 0 | 0 | 0 | 0 | 0 | 0 | 0 | 0 | 2 |
| 6 | ALCCA S | 16 | 32 | 14 | 17 | 16 | 20 | 17 | 11 | 7 | 29 | 41 | 23 | 10 | 12 | 19 | 22 | 21 | 27 | 17 | 48 |
| 7 | ALCCH<br>R | 11 | 10 | 10 | 4 | 8 | 16 | 13 | 6 | 6 | 29 | 22 | 23 | 11 | 14 | 21 | 27 | 24 | 16 | 16 | 48 |
| 8 | ALCCI<br>N | 16 | 50 | 28 | 10 | 24 | 32 | 16 | 16 | 6 | 42 | 50 | 30 | 26 | 7 | 25 | 18 | 26 | 23 | 7 | 70 |
| 9 | BLYPY<br>R | 1 | 0 | 0 | 0 | 0 | 0 | 0 | 0 | 1 | 0 | 0 | 0 | 0 | 0 | 0 | 0 | 0 | 1 | 0 | 0 |
| 10 | BRALE<br>U | 4 | 1 | 0 | 2 | 4 | 1 | 5 | 2 | 4 | 3 | 3 | 2 | 7 | 1 | 2 | 5 | 7 | 3 | 5 | 5 |
| 11 | BRAM<br>ON | 0 | 0 | 0 | 0 | 0 | 0 | 0 | 1 | 0 | 2 | 0 | 1 | 0 | 0 | 0 | 0 | 0 | 0 | 0 | 1 |
| 12 | CERDI<br>S | 0 | 1 | 2 | 0 | 2 | 2 | 0 | 1 | 0 | 1 | 0 | 0 | 0 | 0 | 0 | 0 | 0 | 0 | 0 | 0 |
| 13 | CETFO<br>R | 0 | 1 | 0 | 1 | 1 | 4 | 0 | 1 | 1 | 3 | 1 | 0 | 1 | 2 | 2 | 2 | 1 | 4 | 2 | 2 |
| 14 | CISCHI | 0 | 0 | 1 | 0 | 0 | 0 | 0 | 0 | 0 | 0 | 0 | 1 | 0 | 0 | 0 | 0 | 0 | 0 | 0 | 0 |
| 15 | DENFR<br>O | 0 | 0 | 0 | 0 | 0 | 0 | 0 | 0 | 0 | 0 | 0 | 0 | 0 | 0 | 0 | 0 | 0 | 0 | 0 | 1 |
| 16 | ELAFO<br>R | 0 | 0 | 0 | 0 | 0 | 0 | 0 | 0 | 0 | 0 | 0 | 0 | 0 | 0 | 0 | 0 | 0 | 0 | 0 | 1 |
| 17 | FICHY<br>P | 18 | 13 | 8 | 13 | 11 | 14 | 10 | 7 | 6 | 18 | 19 | 13 | 4 | 2 | 15 | 10 | 10 | 16 | 4 | 12 |
| 18 | FICMO<br>N | 2 | 3 | 5 | 3 | 5 | 6 | 6 | 1 | 2 | 7 | 1 | 0 | 0 | 2 | 6 | 2 | 4 | 3 | 3 | 9 |
| 19 | FICWE<br>S | 0 | 0 | 0 | 0 | 0 | 0 | 0 | 0 | 0 | 0 | 1 | 0 | 0 | 0 | 0 | 0 | 0 | 0 | 0 | 0 |
| 20 | GARC<br>AE | 3 | 2 | 1 | 1 | 3 | 3 | 5 | 3 | 1 | 11 | 4 | 6 | 1 | 0 | 2 | 4 | 3 | 4 | 1 | 2 |
| 21 | GARER<br>Y | 3 | 7 | 6 | 2 | 2 | 5 | 2 | 2 | 6 | 8 | 4 | 4 | 3 | 4 | 2 | 6 | 2 | 8 | 4 | 11 |
| 22 | GARSQ<br>U | 0 | 0 | 1 | 0 | 0 | 0 | 0 | 0 | 0 | 1 | 0 | 0 | 0 | 0 | 4 | 0 | 2 | 3 | 0 | 0 |
| 23 | GARSU<br>B | 1 | 0 | 0 | 0 | 0 | 0 | 0 | 0 | 0 | 0 | 0 | 0 | 0 | 1 | 0 | 0 | 0 | 0 | 0 | 0 |
| 24 | HARER<br>Y | 0 | 2 | 0 | 0 | 1 | 0 | 0 | 0 | 2 | 1 | 1 | 0 | 0 | 0 | 0 | 0 | 0 | 0 | 0 | 0 |
| 25 | LEIAR<br>G | 0 | 0 | 0 | 0 | 0 | 0 | 0 | 0 | 0 | 0 | 1 | 0 | 0 | 0 | 1 | 2 | 0 | 0 | 1 | 0 |
| 26 | LEILUT | 0 | 0 | 0 | 0 | 0 | 0 | 0 | 2 | 0 | 0 | 0 | 0 | 1 | 0 | 0 | 2 | 0 | 0 | 0 | 0 |
| 27 | LIObU<br>G | 0 | 0 | 0 | 0 | 1 | 0 | 0 | 0 | 0 | 0 | 0 | 0 | 0 | 0 | 0 | 2 | 0 | 0 | 0 | 0 |
| 28 | MUSFE<br>R | 0 | 0 | 0 | 0 | 0 | 0 | 0 | 0 | 0 | 0 | 2 | 2 | 0 | 0 | 1 | 0 | 0 | 0 | 0 | 0 |
| 29 | MUSH<br>OD | 1 | 0 | 0 | 3 | 0 | 0 | 0 | 0 | 0 | 0 | 0 | 0 | 0 | 0 | 0 | 0 | 0 | 0 | 0 | 0 |
| 30 | MYILE<br>U | 1 | 3 | 5 | 1 | 6 | 6 | 5 | 4 | 5 | 8 | 3 | 4 | 2 | 1 | 5 | 5 | 9 | 9 | 9 | 6 |
| 31 | NILGR<br>A | 7 | 10 | 8 | 3 | 7 | 4 | 6 | 7 | 5 | 13 | 5 | 8 | 3 | 1 | 3 | 5 | 4 | 7 | 2 | 10 |
| 32 | NILMA<br>C | 0 | 1 | 0 | 0 | 0 | 0 | 0 | 0 | 0 | 1 | 1 | 5 | 0 | 0 | 0 | 0 | 1 | 0 | 0 | 6 |
| 33 | NILSU<br>N | 16 | 1 | 10 | 4 | 11 | 12 | 16 | 4 | 2 | 17 | 11 | 0 | 4 | 1 | 6 | 6 | 5 | 4 | 1 | 2 |
| 34 | ORTCU<br>C | 0 | 2 | 0 | 0 | 1 | 0 | 4 | 0 | 0 | 0 | 0 | 4 | 1 | 3 | 3 | 2 | 4 | 3 | 0 | 0 |
| 35 | PARNI<br>P | 17 | 33 | 14 | 8 | 7 | 17 | 7 | 6 | 11 | 26 | 33 | 7 | 8 | 1 | 16 | 18 | 32 | 19 | 13 | 26 |
| 36 | PARRU<br>F | 0 | 0 | 9 | 1 | 2 | 5 | 2 | 0 | 2 | 0 | 5 | 1 | 0 | 0 | 1 | 0 | 1 | 1 | 0 | 4 |
| 37 | PARSPI | 2 | 0 | 5 | 0 | 1 | 0 | 1 | 0 | 2 | 1 | 0 | 0 | 1 | 0 | 0 | 1 | 0 | 0 | 0 | 3 |
| 38 | PHYCH<br>L | 0 | 1 | 0 | 0 | 0 | 0 | 0 | 0 | 0 | 1 | 3 | 0 | 0 | 0 | 0 | 0 | 0 | 1 | 0 | 1 |
| 39 | PHYRE<br>G | 7 | 7 | 6 | 4 | 3 | 2 | 9 | 2 | 1 | 14 | 6 | 9 | 1 | 3 | 0 | 9 | 10 | 5 | 2 | 9 |
| 40 | PITNIP | 0 | 0 | 0 | 0 | 1 | 0 | 0 | 0 | 0 | 0 | 0 | 0 | 0 | 0 | 0 | 0 | 0 | 0 | 0 | 0 |
| 41 | PN0PU<br>S | 0 | 0 | 0 | 0 | 0 | 0 | 0 | 0 | 0 | 2 | 0 | 0 | 0 | 0 | 0 | 0 | 0 | 0 | 0 | 0 |
| 42 | PNOPU<br>S | 5 | 2 | 2 | 0 | 1 | 0 | 1 | 0 | 1 | 2 | 1 | 1 | 0 | 0 | 0 | 1 | 1 | 0 | 0 | 1 |
| 43 | POMFE<br>R | 0 | 1 | 2 | 1 | 0 | 3 | 1 | 4 | 0 | 2 | 1 | 2 | 2 | 2 | 3 | 0 | 1 | 0 | 0 | 1 |
| 44 | POMR<br>UF | 3 | 4 | 0 | 2 | 2 | 1 | 3 | 2 | 1 | 2 | 6 | 3 | 0 | 1 | 4 | 3 | 8 | 5 | 2 | 7 |
| 45 | PSADA<br>L | 0 | 0 | 0 | 0 | 0 | 0 | 0 | 1 | 0 | 1 | 0 | 0 | 0 | 0 | 0 | 0 | 0 | 0 | 0 | 0 |
| 46 | RHIAL<br>B | 2 | 2 | 2 | 1 | 0 | 4 | 3 | 2 | 2 | 9 | 8 | 3 | 3 | 1 | 0 | 1 | 5 | 5 | 4 | 3 |
| 47 | RIMM<br>AL | 2 | 0 | 0 | 0 | 2 | 1 | 0 | 0 | 0 | 0 | 0 | 0 | 1 | 0 | 1 | 0 | 0 | 0 | 0 | 0 |
| 48 | SASOC<br>H | 0 | 0 | 0 | 0 | 0 | 2 | 1 | 2 | 1 | 2 | 0 | 1 | 1 | 0 | 1 | 3 | 2 | 2 | 2 | 6 |
| 49 | SEIAFF | 9 | 8 | 8 | 2 | 6 | 6 | 7 | 5 | 6 | 15 | 12 | 9 | 3 | 2 | 6 | 6 | 4 | 4 | 2 | 13 |
| 50 | SEIPOL | 10 | 13 | 4 | 7 | 4 | 11 | 11 | 4 | 8 | 24 | 10 | 9 | 4 | 7 | 5 | 14 | 8 | 5 | 3 | 22 |
| 51 | SPECA<br>U | 3 | 2 | 1 | 1 | 7 | 5 | 4 | 2 | 1 | 7 | 5 | 1 | 0 | 0 | 2 | 0 | 3 | 3 | 2 | 3 |
| 52 | SPHHU<br>M | 1 | 0 | 0 | 0 | 0 | 0 | 0 | 0 | 0 | 0 | 0 | 1 | 0 | 0 | 0 | 0 | 0 | 0 | 0 | 0 |
| 53 | STACH<br>R | 8 | 15 | 10 | 4 | 6 | 14 | 9 | 8 | 3 | 19 | 9 | 8 | 7 | 7 | 16 | 15 | 14 | 11 | 10 | 19 |
| 54 | STANI<br>G | 0 | 0 | 0 | 0 | 0 | 0 | 0 | 0 | 0 | 0 | 0 | 1 | 0 | 0 | 0 | 0 | 0 | 0 | 0 | 0 |
| 55 | STARU<br>F | 12 | 16 | 12 | 10 | 8 | 16 | 16 | 7 | 7 | 19 | 33 | 31 | 17 | 16 | 38 | 31 | 26 | 29 | 13 | 37 |
| 56 | TESCA<br>S | 1 | 0 | 2 | 3 | 0 | 0 | 0 | 0 | 0 | 3 | 5 | 6 | 0 | 1 | 0 | 0 | 0 | 0 | 0 | 8 |
| 57 | TESCY<br>A | 4 | 4 | 2 | 1 | 3 | 2 | 3 | 3 | 0 | 1 | 11 | 3 | 2 | 2 | 4 | 2 | 1 | 1 | 0 | 6 |
| 58 | TESOLI | 2 | 1 | 2 | 0 | 3 | 3 | 2 | 3 | 1 | 2 | 3 | 2 | 0 | 2 | 2 | 3 | 0 | 1 | 1 | 5 |
| 59 | TICHO<br>D | 0 | 1 | 0 | 2 | 0 | 0 | 0 | 0 | 0 | 3 | 4 | 3 | 1 | 1 | 2 | 2 | 1 | 1 | 0 | 6 |
| 60 | XIPSUP | 0 | 0 | 2 | 0 | 2 | 1 | 0 | 2 | 2 | 5 | 3 | 5 | 0 | 0 | 1 | 0 | 1 | 1 | 2 | 5 |
| 61 | YUHFL<br>A | 12 | 0 | 0 | 1 | 0 | 1 | 2 | 2 | 2 | 9 | 6 | 3 | 8 | 2 | 2 | 3 | 9 | 1 | 1 | 14 |

**Supp. Table S3:** Similarity percentage (SIMPER) routines of species that contribute to 50% cumulative dissimilarity between the two habitats. Note: *Schoeniparus cinereus* and *Tesia castaneocoronata* have not been included in Figure 2 (NMDS)

| Species | Species code | average | sd | ratio | ava | avb | cumsu<br>m | p<br>value | significanc<br>e |
| --- | --- | --- | --- | --- | --- | --- | --- | --- | --- |
| *Niltava sundara* | NILSUN | 0.009681 | 0.006145 | 1.5754 | 0.2031 | 0.1182 | 0.038 | 0.008 | ** |
| *Stachyris ruficeps* | STARUF | 0.009032 | 0.004001 | 2.2573 | 0.249 | 0.3419 | 0.074 | 0.001 | *** |
| *Yuhina flavicollis* | YUHFLA | 0.007835 | 0.005886 | 1.3311 | 0.0912 | 0.1336 | 0.105 | 0.227 |  |
| *Lioparus chrysotis* | ALCCHR | 0.007678 | 0.004217 | 1.8208 | 0.2292 | 0.3063 | 0.136 | 0.001 | *** |
| *Phyllergates*<br>*cucullatus* | ORTCUC | 0.007159 | 0.005321 | 1.3454 | 0.0304 | 0.0818 | 0.164 | 0.048 | * |
| *Psittiparus ruficeps* | PARRUF | 0.006938 | 0.005376 | 1.2905 | 0.0792 | 0.0472 | 0.192 | 0.239 |  |
| *Schoeniparus*<br>*cinereus* | ALCCIN | 0.00692 | 0.004932 | 1.4032 | 0.3324 | 0.3265 | 0.219 | 0.879 |  |
| *Tickellia hodgsoni* | TICHOD | 0.006553 | 0.003706 | 1.7682 | 0.0277 | 0.0825 | 0.245 | 0.005 | ** |
| *Suthora nipalensis* | PARNIP | 0.00652 | 0.005306 | 1.2288 | 0.2616 | 0.2528 | 0.271 | 0.944 |  |
| *Spelaeoris caudatus* | SPECAU | 0.006052 | 0.004935 | 1.2263 | 0.1216 | 0.07 | 0.295 | 0.033 | * |
| *Abroscopus schisticeps* | ABRSCH | 0.005949 | 0.005781 | 1.029 | 0.1275 | 0.1021 | 0.318 | 0.685 |  |
| *Tesia castaneocoronata* | TESCAS | 0.005933 | 0.005626 | 1.0546 | 0.0413 | 0.0508 | 0.342 | 0.811 |  |
| *Pomatorhinus superciliaris* | XIPSUP | 0.005774 | 0.004417 | 1.3074 | 0.0651 | 0.0655 | 0.365 | 0.842 |  |
| *Pomatorhinus ferruginosus* | POMFER | 0.00576 | 0.004343 | 1.3263 | 0.0688 | 0.0625 | 0.388 | 0.922 |  |
| *Brachypteryx leucophrys* | BRALEU | 0.005728 | 0.004386 | 1.3059 | 0.1109 | 0.1312 | 0.41 | 0.433 |  |
| *Anthipes monileger* | FICMON | 0.005618 | 0.004838 | 1.1611 | 0.1383 | 0.0966 | 0.433 | 0.121 |  |
| *Machlolophus spilonotus* | PARSPI | 0.005486 | 0.004884 | 1.1234 | 0.0589 | 0.023 | 0.455 | 0.183 |  |
| *Sasia ochracea* | SASOCH | 0.005414 | 0.004066 | 1.3315 | 0.0459 | 0.0739 | 0.476 | 0.234 |  |
| *Ficedula hyperythra* | FICHYP | 0.005386 | 0.004097 | 1.3149 | 0.245 | 0.2015 | 0.497 | 0.038 | * |
| *Niltava grandis* | NILGRA | 0.005301 | 0.003144 | 1.6863 | 0.1875 | 0.1365 | 0.518 | 0.001 | *** |

Significance codes: '\*\*\*' = $p < 0.001$ , '\*\*' = $p < 0.01$ , '\*' = $p < 0.05$
average = average contribution of the species to between-group dissimilarity
ava, avb = average abundances of each habitat (primary and logged)

**Figure S1:**
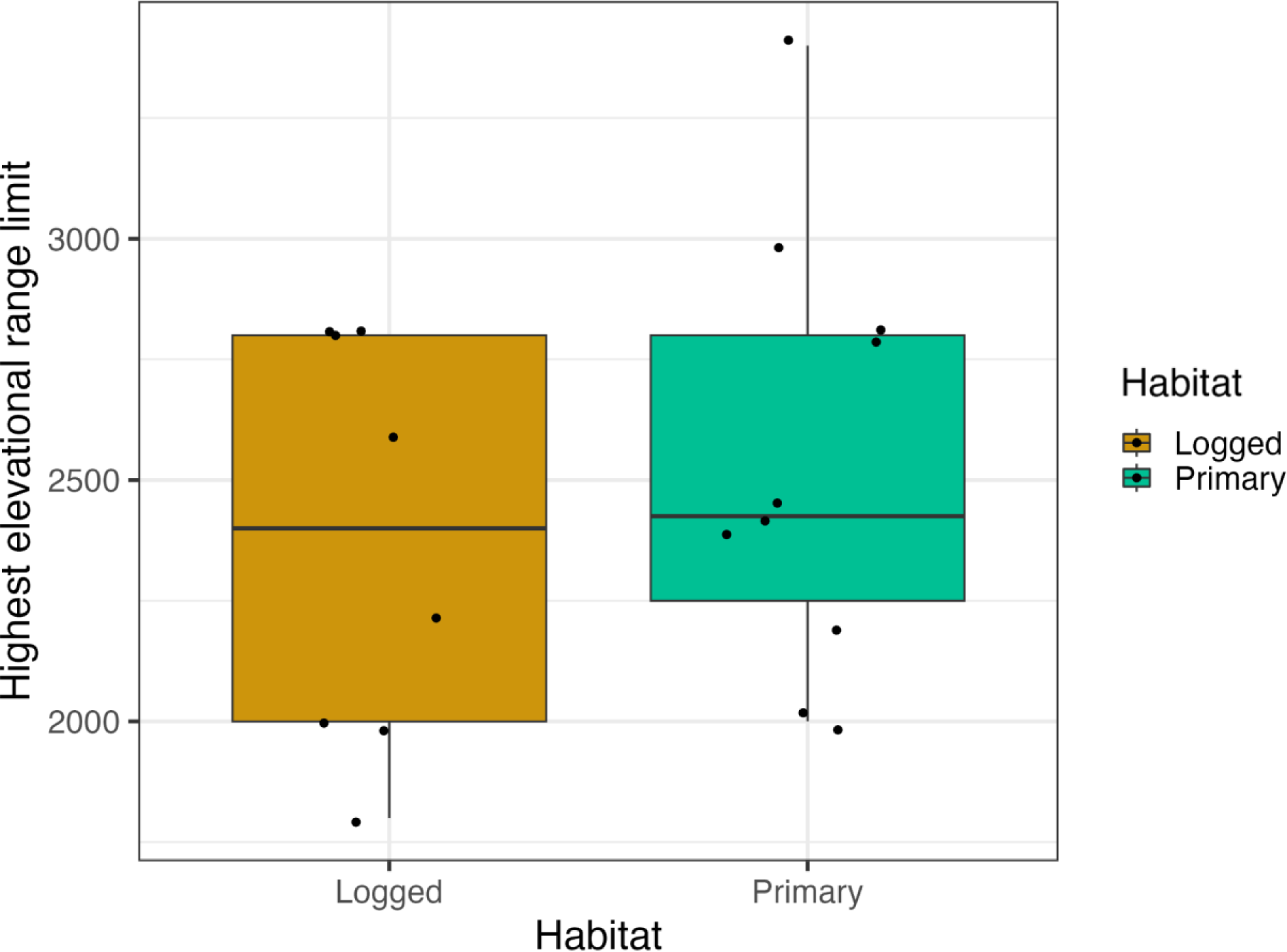
Highest breeding elevation range of the species that contributed to 50% dissimilarity in each habitat. The box shows the middle 50% of the data, with the horizontal line inside the box indicating the median value. The whiskers of the boxplot extend from the box to show the minimum and maximum values of the data. Each dot represents individual data points and is plotted with a jitter to avoid overlap.

**Figure S2:**
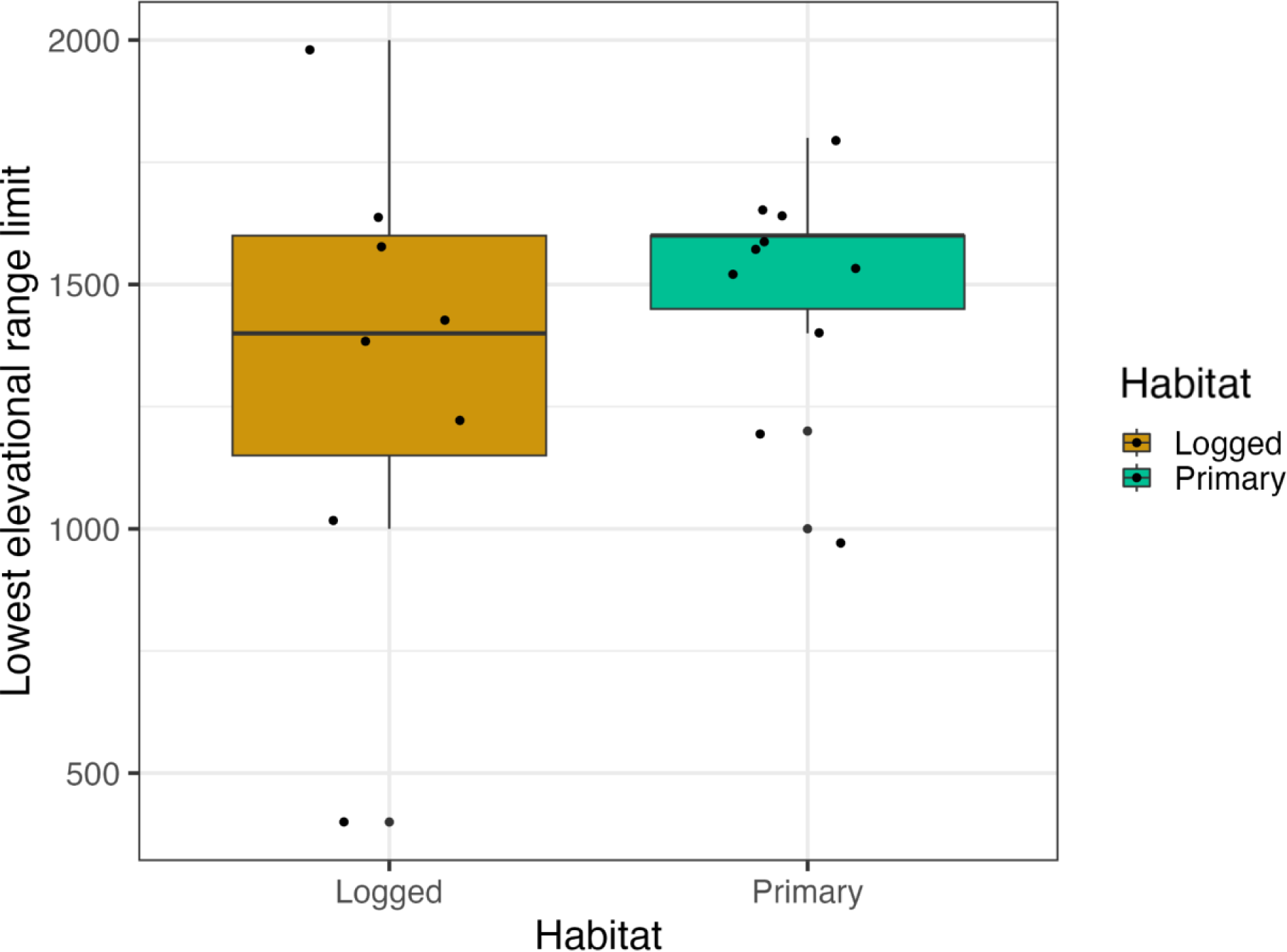
Lowest breeding elevation range of the species that contributed to 50% dissimilarity in each habitat. The box shows the middle 50% of the data, with the horizontal line inside the box indicating the median value. The whiskers of the boxplot extend from the box to show the minimum and maximum values of the data. Each dot represents individual data points and is plotted with a jitter to avoid overlap.

**Figure S3:**
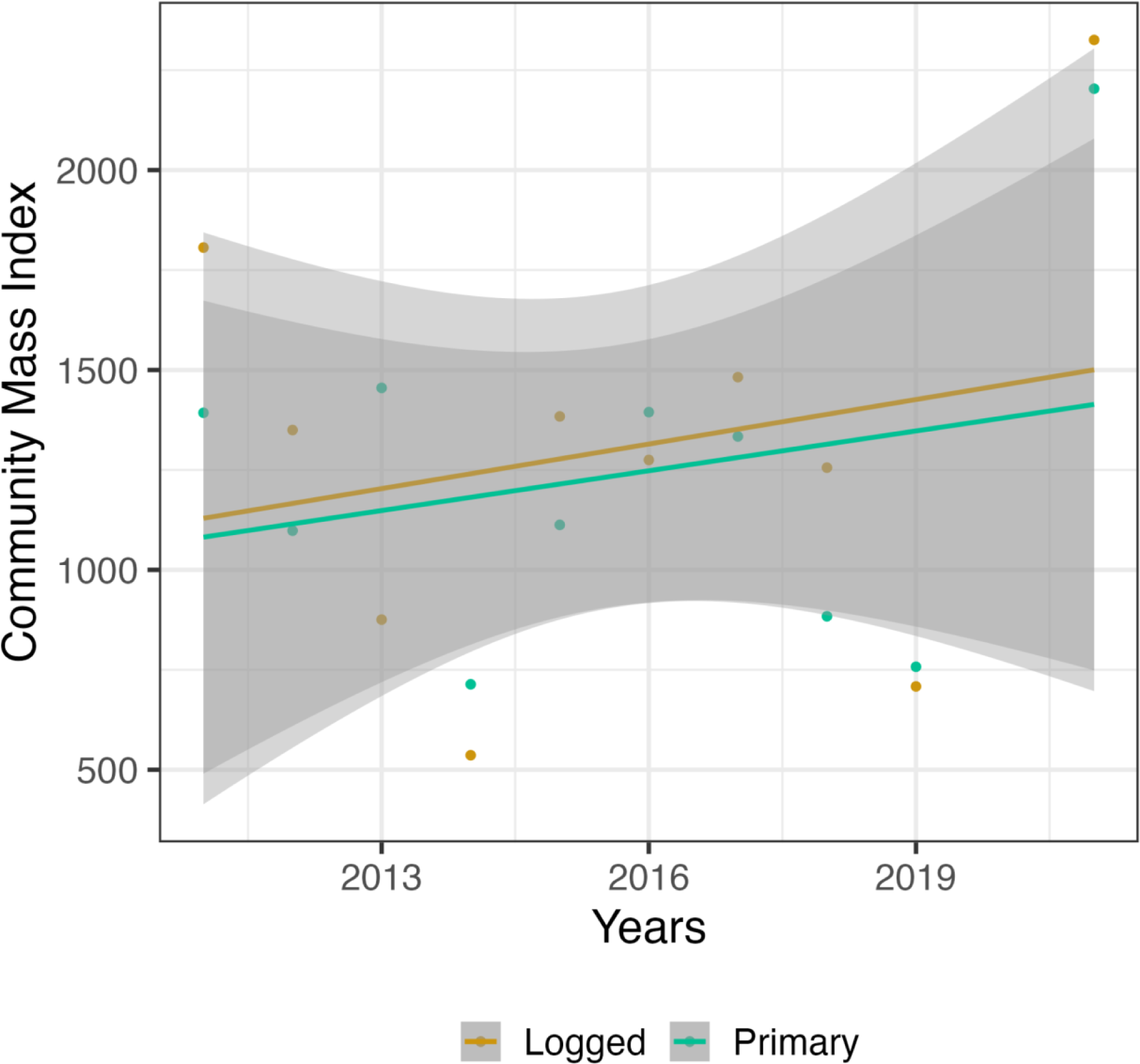
Abundance weighted mass index over time of the species that contributed to 50% dissimilarity in each habitat.

**Figure S4:**
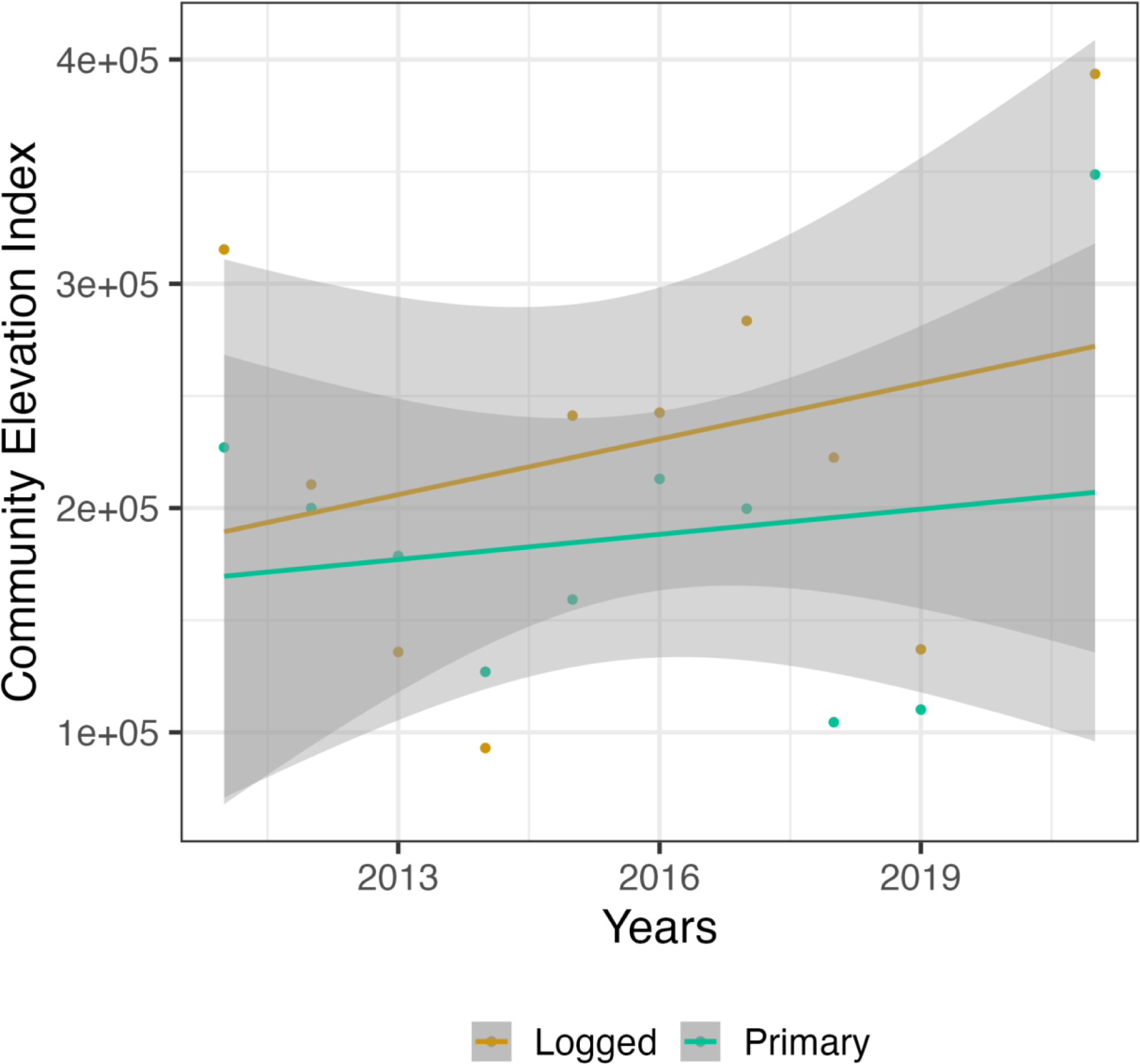
Abundance weighted mean breeding elevation index over time of the species that contributed to 50% dissimilarity in each habitat.

## Notes

### Competing Interest Statement

The authors have declared no competing interest.

### Summary of Updates

Some new analyses have been added

